# The tubule-sheet continuum model links the pattern of nuclear envelope assembly to nuclear envelope integrity

**DOI:** 10.1101/2024.07.14.603427

**Authors:** Sanjana Arun, Gengjing Zhao

## Abstract

Nuclear envelope (NE) fragility is associated with a variety of human diseases. However, major questions about the causes of NE fragility are unresolved. In particular, the relationship between NE fragility and NE assembly is unclear. Here, using a panel of 13 mammalian cell lines, we show that normal cells and some cancer cells assemble the NE using distinct patterns from opposite ends of the “tubule-sheet continuum”, and that the NE assembly pattern more utilised in cancer cells is a source of NE fragility. In normal cells, the mitotic endoplasmic reticulum (ER) morphology is dominated by ER tubules or small sheets and the NE assembles by “membrane infiltration”. By contrast, in some cancer cells, the mitotic ER morphology is dominated by large ER sheets and the NE assembles by “lateral sheet expansion”. Experimental manipulations that promote mitotic ER sheet formation and NE assembly by lateral sheet expansion in normal cells result in NE fragility. Thus, we propose that NE assembly by lateral sheet expansion is a source of NE fragility in cancer cells.

## Introduction

The nuclear envelope (NE) is an important physical barrier that protects the genome and is frequently disrupted in human diseases. First identified in fibroblasts from laminopathy patients (De Vos et al., 2011; Vigouroux et al., 2001), it is now known that fragile NEs that undergo loss of integrity or “ruptures” are also a common feature of cancer cells (Coy et al., 2023; Vargas et al., 2012). The causes of NE ruptures in cancer cells include mechanical stress from nuclear confinement or cell migration through constrictions (Denais et al., 2016; Hatch and Hetzer, 2016; Irianto et al., 2017; Jung-Garcia et al., 2023; Nader et al., 2021; Raab et al., 2016; Takaki et al., 2017). Defects from NE assembly are another source of NE ruptures, as occurs on micronuclei and chromosome bridges—abnormal nuclear structures common in cancer cells, and can result in chromosome fragmentation, transcription and chromatin defects, and innate immune activation (Crasta et al., 2012; Harding et al., 2017; Hatch et al., 2013; Maciejowski et al., 2015; Mackenzie et al., 2017; Mohr et al., 2021; Papathanasiou et al., 2023; Tang et al., 2022; Zhang et al., 2015).

Given its important role as a physical barrier, it is perhaps counter-intuitive that in metazoan cells, the NE completely breaks down each cell division and reassembles from the mitotic endoplasmic reticulum (ER) at mitotic exit (Deolal et al., 2024; Ungricht and Kutay, 2017). During reassembly, two groups of NE proteins, the “core” (e.g., the Barrier-to-autointegration factor, BAF) and “non-core” (e.g., the Nuclear Pore Complexes, NPCs) proteins, assemble on chromatin concomitantly with NE membranes (Dechat et al., 2004; Haraguchi et al., 2008; Haraguchi et al., 2000; Haraguchi et al., 2001; Kutay et al., 2021; LaJoie and Ullman, 2017; Liu et al., 2018; Liu and Pellman, 2020; Lu et al., 2011; Olmos et al., 2015; Otsuka et al., 2018; Samwer et al., 2017; Vietri et al., 2015; von Appen et al., 2020; Zhao et al., 2023). In HeLa cells, the core and non-core NE proteins transiently form two distinct NE subdomains around chromatin: the core NE proteins accumulate in the spindle-adjacent core chromatin regions (Dechat et al., 2004; Haraguchi et al., 2008; Haraguchi et al., 2000; Haraguchi et al., 2001; Liu et al., 2018; Samwer et al., 2017; von Appen et al., 2020; Zhao et al., 2023), whereas the non-core NE proteins assemble on the peripheral non-core chromatin regions away from the spindle, and are depleted from the core chromatin regions (Dechat et al., 2004; Haraguchi et al., 2008; Haraguchi et al., 2000; Liu et al., 2018; Lu et al., 2011; Otsuka et al., 2018; Zhao et al., 2023). However, in other cell types, both core and non-core NE proteins assemble in the core chromatin regions and there is not a distinct separation of the two NE subdomains (Zhao et al., 2023), suggesting cell type differences in NE assembly. Importantly, failure to assemble both groups of NE proteins results in NE fragility, as occurs on micronuclei and chromosome bridges, which primarily assemble core but are highly deficient in non-core proteins (de Castro et al., 2018; Hatch et al., 2013; Liu et al., 2018; Zhao et al., 2023).

We previously identified two distinct NE assembly patterns in human cells with different mitotic ER morphologies that explain the cell type differences in NE subdomain formation (Zhao et al., 2023). In cell types whose mitotic ER morphology is dominated by large ER sheets, the NE first assembles on the peripheral non-core chromatin regions and then extends into the spindle-adjacent core chromatin regions (“lateral sheet expansion”; (Lu et al., 2011; Otsuka and Ellenberg, 2018; Otsuka et al., 2018)). This NE assembly pattern results in the formation of distinct core and non-core NE subdomains, as NPCs do not assemble on the NE extensions from large ER sheets. By contrast, in cell types whose mitotic ER morphology is dominated by ER tubules or small sheets, NE membranes and NPCs are recruited simultaneously to the core and non-core chromatin regions by mitotic actin filaments (“membrane infiltration”), and therefore distinct NE subdomains are not formed. These two NE assembly patterns define the opposite ends of the “tubule-sheet continuum”, which we proposed allows cell types with any mix of ER tubules and ER sheets to efficiently assemble the NE using one or a combination of these two NE assembly patterns. However, how diverse cell types distribute on the tubule-sheet continuum, and whether the NE assembly patterns have distinct consequences on nuclear integrity, are unclear.

Here, we extend our analysis of the tubule-sheet continuum model to 13 mammalian cell lines. We show that all 9 non-transformed cell lines examined (hereafter referred to simply as “normal”) lie on the “ER tubules or small sheets” end of the continuum and assemble the NE by membrane infiltration. By contrast, 3 out of the 4 cancer cell lines examined are biased towards the “large ER sheets” end of the continuum and assemble the NE by lateral sheet expansion. Additionally, we show that shifting the tubule-sheet balance in normal cells towards large ER sheets and the NE assembly pattern towards lateral sheet expansion results in NE fragility. Thus, we propose that NE assembly by lateral sheet expansion is a source of NE fragility in cancer cells.

## Results

### The mitotic ER morphology is dominated by ER tubules or small sheets in normal cells and by large ER sheets in some cancer cells

We determined the mitotic ER morphology in 10 mammalian cell lines in addition to the three we assessed previously: U2OS, HeLa, and RPE-1 (Zhao et al., 2023). Our new analysis includes eight normal cell lines: six hTERT-immortalised primary human cell lines, and the rat NRK-52E (de Larco and Todaro, 1978) and mouse NIH-3T3 (Jainchill et al., 1969) lines, and two additional human cancer cell lines: Huh-7 and HCT116 (**Table S1**).

We assessed the mitotic ER morphology in anaphase or telophase cells by confocal live-cell microscopy. Using the ER membrane marker GFP-Sec61β, we first verified our prior findings (Zhao et al., 2023) that the mitotic ER in HeLa consists of large ER sheets (median ER length ∼1300 nm), whereas the mitotic ER in U2OS consists of ER tubules or small sheets (median ER length ∼650 nm; **Fig. 1, A and B**). The measured median ER lengths in HeLa or U2OS cells were consistent regardless of the live-cell ER marker used (GFP-Sec61β (this study), YFP-STIM1 or ER-trackerRed (Zhao et al., 2023)), demonstrating that these markers can be used interchangeably to determine mitotic ER morphology.

**Figure 1.**
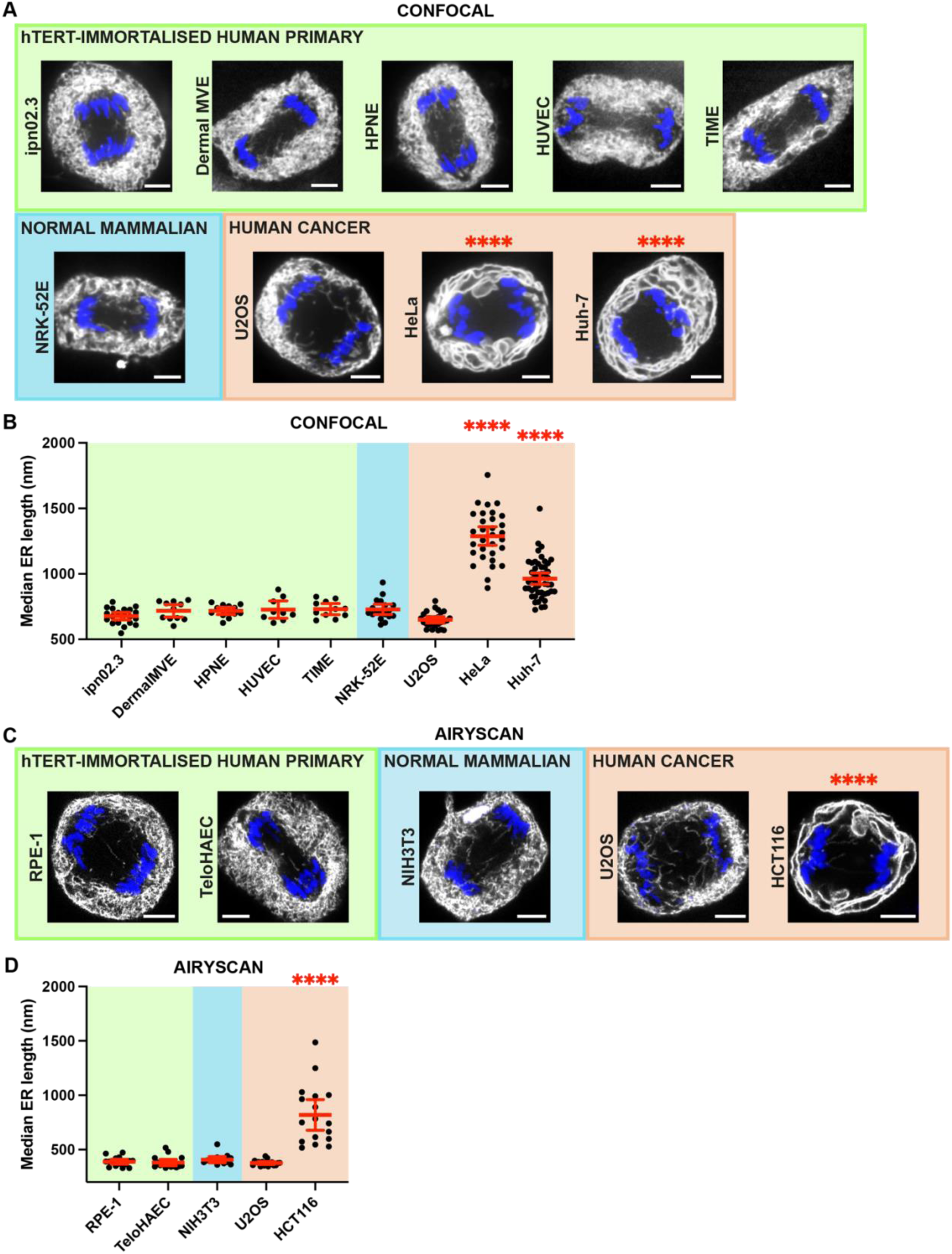
The mitotic ER morphology in 13 mammalian cell lines. **(A–D)** The mitotic ER is dominated by ER tubules or small sheets in the 9 normal cell lines (hTERT-immortalised human primary and normal mammalian), and by large ER sheets in 3 out of the 4 human cancer cell lines examined. **(A)** Live-cell confocal images of the ER in the indicated anaphase or telophase cells. Note the large mitotic ER sheets (asterisks) in HeLa or Huh-7 (cancer) cells. **(B)** Median ER lengths measured from live-cell confocal images as shown in (A). **(C and D)** Same as in (A and B) except for Airyscan images. Note the large mitotic ER sheets (asterisks) in HCT116 (cancer) cells. Images: single focal plane; ER (GFP-Sec61β, YFP-STIM1, or ER-trackerRed), white; DNA (SiR-DNA), blue; scale bars: 5 μm. Sample size: N = 20, 11, 13, 9, 11, 18, 26, 31, 48 from left to right in (B); N = 17, 16, 16, 13, 17 from left to right in (D). Quantification: ^∗∗∗∗^p < 0.0001, or not significant (not labelled), Kruskal-Wallis tests, comparisons to ipn02.3 in (B) or RPE-1 in (D); error bars: mean with 95% CI.

Surprisingly, the mitotic ER is dominated by ER tubules or small sheets in all six normal cell lines we examined by confocal live-cell imaging (median ER lengths between ∼630 nm and ∼730 nm; **Fig. 1, A and B**). As we cannot resolve structures below the diffraction limit, we likely overestimated the lengths of ER tubules. Therefore, we assessed the mitotic ER morphology in three other normal cell lines (RPE-1, TeloHAEC, and NIH-3T3) by Airyscan live-cell microscopy (**Fig. 1 C**). As expected, the measured median ER length was shorter (∼380 nm) in telophase U2OS cells from the Airyscan as compared to the confocal images. Additionally, the median ER lengths in RPE-1, TeloHAEC, and NIH-3T3 cells were similar to that of U2OS cells (**Fig. 1 D**). These results confirm our previous findings in RPE-1 cells (Zhao et al., 2023), and demonstrate that the mitotic ER morphology is dominated by ER tubules or small sheets in all 9 normal mammalian cell lines examined. By contrast, large ER sheets were more abundant in the cancer cell lines HeLa, Huh-7, and HCT116 as compared to the normal cells. As in HeLa cells, the mitotic ER in Huh-7 and HCT116 cells is dominated by large ER sheets (**Fig. 1, A and C)** and had significantly longer median ER lengths than normal cells (**Fig. 1, B and D**). The reticulated ER morphology in U2OS (cancer) and the intermediate ER morphology in Huh-7 (in between HeLa and U2OS) suggest that cancer cells adopt a range of mitotic ER morphologies, but are biased towards large ER sheets.

Interestingly, prior work reported similar mitotic ER lengths in Huh-7 (cancer) and NRK-52E (normal) cells (Puhka et al., 2012). To address the discrepancy with our results (**Fig. 1, A and B**), we measured the density of mitotic F-actin in the subcortical cytoplasm of telophase cells, which is required for maintaining ER tubules or small sheets (Zhao et al., 2023), as an orthogonal readout of the mitotic ER morphology in these two cell lines. We found that the density of mitotic F-actin in the subcortical cytoplasm is significantly higher in NRK-52E as compared to Huh-7 cells, while the density of cortical F-actin is similar in the two cell lines (**Fig. S1, A and B**). Combined with our previous findings, these results show that the mitotic ER morphology is dominated by ER tubules or small sheets in cells with high mitotic F-actin density in the subcortical cytoplasm (NRK-52E (this study), RPE-1 or U2OS (Zhao et al., 2023)). By contrast, the mitotic ER morphology is dominated by large ER sheets in cells with low mitotic F-actin density in the subcortical cytoplasm (Huh-7 (this study) or HeLa (Zhao et al., 2023)).

In summary, our findings suggest that the mitotic ER morphology in all normal cells examined is dominated by ER tubules or small sheets, whereas the mitotic ER morphology in some cancer cells is dominated by large ER sheets.

### The NE assembles by membrane infiltration in normal cells and lateral sheet expansion in some cancer cells

As the mitotic ER morphology controls the pattern of NE assembly (Zhao et al., 2023), we would expect the differences in mitotic ER morphology that we observed in the different cell lines to be reflected in their NE assembly patterns. Therefore, we followed NE assembly in the different cell lines by time-lapse imaging (**Fig. 2 A and Fig. S2**), and determined the ratio of NE recruitment in the inner-core relative to the non-core chromatin region, hereafter referred to as the “NE recruitment ratio”. The NE recruitment ratio is ∼1 for membrane infiltration (for synchronous NE recruitment to the inner-core and non-core NE regions) and <1 for lateral sheet expansion (for delayed NE recruitment to the inner-core relative to the non-core NE region; **Fig. 2 B**; (Zhao et al., 2023)). Consistent with having a mitotic ER dominated by ER tubules or small sheets, the NE assembled by membrane infiltration in all normal cell lines examined (**Fig. 2 A and C; and Fig. S2**).

**Figure 2.**
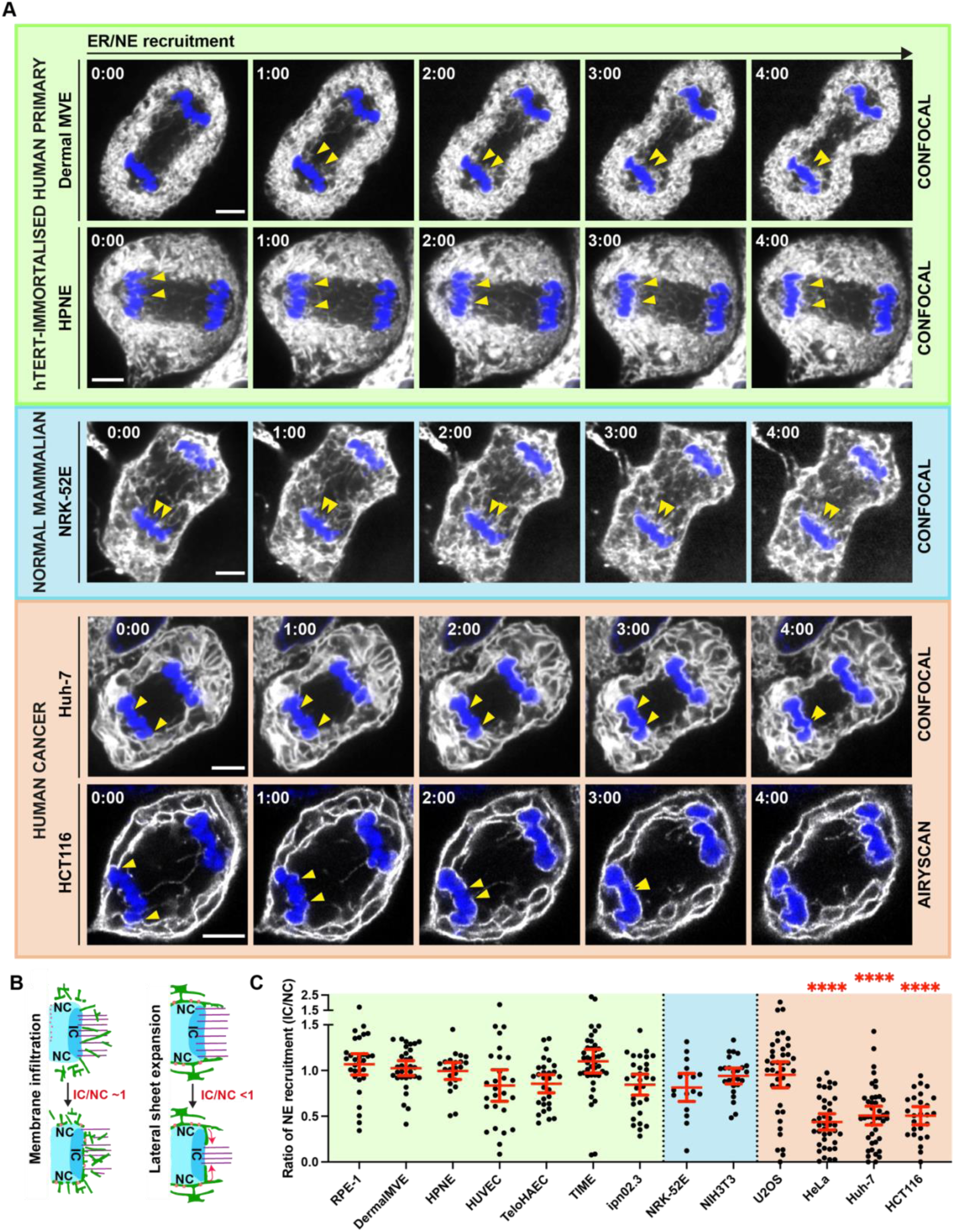
The NE assembly pattern in 13 mammalian cell lines. **(A–C)** The NE assembles by membrane infiltration in the 9 normal cell lines (hTERT-immortalised human primary and normal mammalian), and by lateral sheet expansion in 3 out of the 4 human cancer cell lines examined. Note: images of a subset of cell lines are shown (see Fig. S2 for images of the remaining cell lines). **(A)** Confocal or Airyscan images (as indicated) from time-lapse series showing NE assembly in the indicated cells. Yellow arrowheads: NE recruitment to the IC chromatin region by membrane infiltration in Dermal MVE, HPNE, or NRK-52E (normal) cells; and NE extension into the inner-core (IC) from the non-core (NC) chromatin region by lateral sheet expansion in Huh-7 or HCT116 (cancer) cells. **(B)** Cartoon of the two NE assembly patterns and the expected ratios of NE recruitment in the IC relative to the NC chromatin region (∼1 for membrane infiltration; <1 for lateral sheet expansion). **(C)** Ratio of NE membrane recruitment in the IC relative to the NC in the indicated cells. Shown is the fraction of the IC covered divided by the fraction of the NC covered by ER membranes; ratios are at half-maximal ER membrane recruitment (see Materials and methods). Images: single focal plane; ER (GFP-Sec61β, YFP-STIM1, or ER-trackerRed), white; DNA (SiR-DNA), blue; scale bars: 5 μm. Sample size: N = 30, 32, 22, 27, 29, 41, 30, 17, 25, 40, 39, 42, 26 from left to right in (C). Quantification: ^∗∗∗∗^p < 0.0001, or not significant (not labelled), Kruskal-Wallis tests, comparisons to RPE-1; error bars: mean with 95% CI.

On the other hand, the human cancer cell lines HeLa, Huh-7, and HCT116 have NE recruitment ratios of ∼0.5 (**Fig. 2 C**), suggesting NE assembly by lateral sheet expansion in these cell lines. Timelapse imaging verified the assembly of the NE by lateral sheet expansion from large ER sheets that first wrapped around the periphery of the chromosome mass and then extended into the inner-core chromatin region in these cell lines (**Fig. 2 A and Fig. S2**). Combined with our findings that the mitotic ER is dominated by large ER sheets in HeLa, Huh-7, and HCT116 cells (**Fig. 1**), these data confirm that large ER sheets favour NE assembly by lateral sheet expansion. The membrane infiltration NE assembly pattern in U2OS suggests that cancer cells utilise a range of NE assembly patterns, but are biased towards lateral sheet expansion.

Together, our data suggest that in all normal cells examined, the NE assembles by membrane infiltration from ER tubules or small sheets. By contrast, in some cancer cells, where the mitotic ER is dominated by large ER sheets, the NE assembles by lateral sheet expansion.

### Promoting NE assembly by lateral sheet expansion in RPE-1 cells results in NE regions with protein mis-localisation

Given the distinct NE assembly patterns utilised in the normal cells versus some cancer cells, we assessed the consequences of promoting NE assembly by lateral sheet expansion in normal cells. We used three orthogonal methods (Zhao et al., 2023) to promote mitotic ER sheet formation and hence the lateral sheet expansion NE assembly pattern in RPE-1 cells (**Fig. 3 A**): abolishing interphase ER tubule-microtubule connections (Lu et al., 2009; Terasaki et al., 1986) using Nocodazole (hereafter referred to as “converted”); the knockout of RTN4, a reticulon family protein that generates the high curvature of ER tubules (Voeltz et al., 2006); and the overexpression of the ER sheet-forming protein CLIMP63 (Shibata et al., 2010). We then stained for different core (Emerin, Lamin A, LAP2α, or LAP2β) and non-core (Lamin B1, Lamin B Receptor (LBR), or NPCs) NE proteins and examined their localisation.

**Figure 3.**
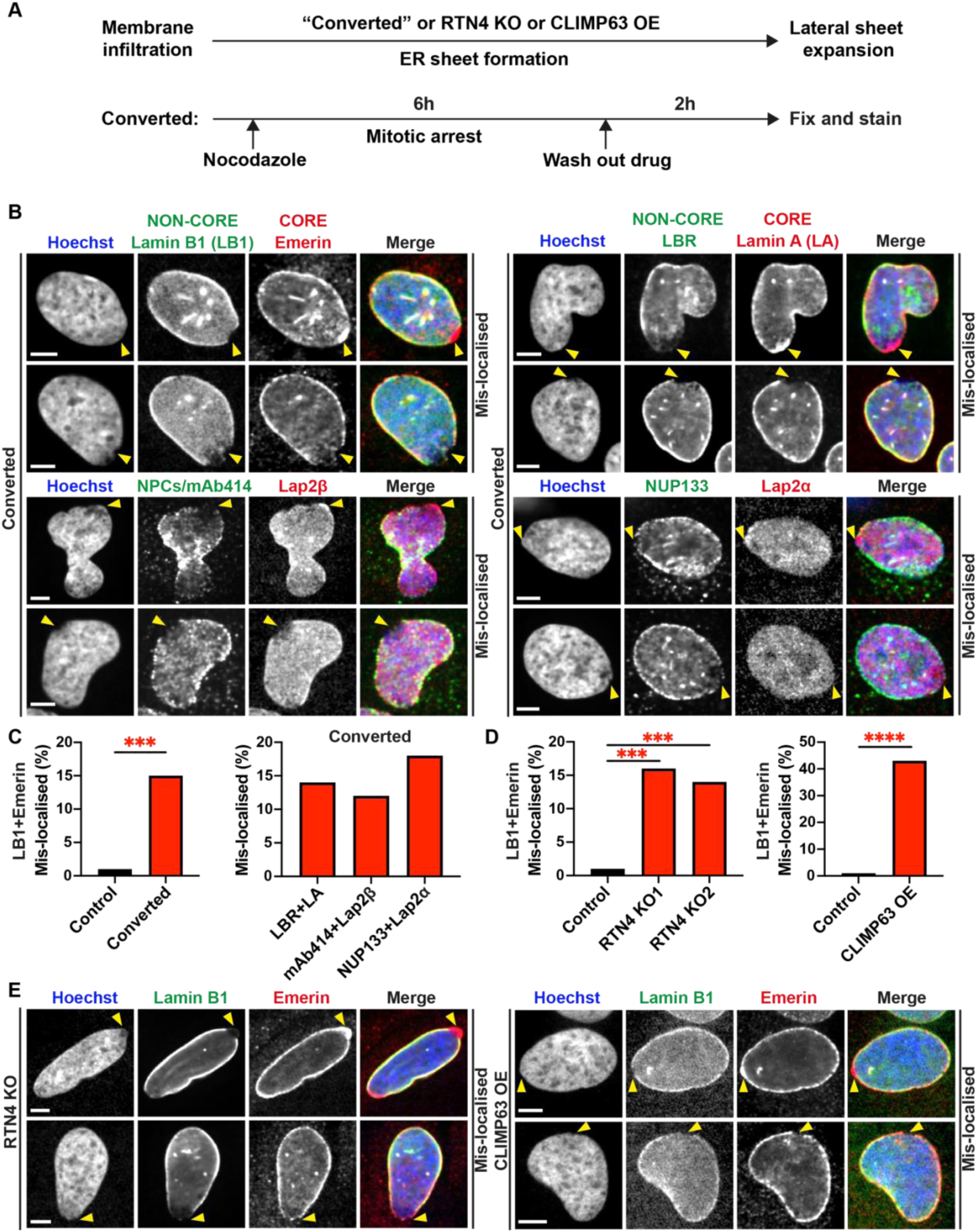
Promoting NE assembly by lateral sheet expansion in RPE-1 cells results in NE regions with protein mis-localisation. **(A)** Experimental scheme. Orthogonal methods of promoting mitotic ER sheet formation and hence the lateral sheet expansion NE assembly pattern (“converted”, RTN4 knockout (KO), or CLIMP63 overexpression (OE); (Zhao et al., 2023)). Converted: depolymerisation of microtubules in interphase and mitosis by Nocodazole-treatment, followed by washing out the drug ((Zhao et al., 2023); Materials and methods). **(B–D)** Mis-localisation of NE proteins in converted, RTN KO or CLIMP63 OE nuclei. **(B)** Co-staining of non-core (Lamin B1 (LB1), Lamin B Receptor (LBR), NPCs/mAb414, or NUP133)) and core (Emerin, Lamin A, Lap2β, or Lap2α) NE markers in converted cells. Arrowheads: NE regions with non-core depletion alongside core enrichment (top panels), or co-depletion of both non-core and core (bottom panels). Note: “mis-localised” refers to either protein mis-localisation event; Lap2α depletion was not observed. **(C)** Left, percentage of control or converted nuclei with NE regions where both Lamin B1 and Emerin (LB1+Emerin) are mis-localised. Control: depolymerisation of microtubules only in mitosis by Nocodazole-treatment, followed by washing out the drug, which does not convert the membrane infiltration NE assembly pattern into lateral sheet expansion ((Zhao et al., 2023); Materials and methods). Right, same as left except for different non-core and core NE markers (as indicated). **(D)** Left, percentage of nuclei with NE regions where both Lamin B1 and Emerin (LB1+Emerin) are mis-localised in control or two RTN4 KO lines (KO1 and KO2). Right, same as left except for CLIMP63 OE cells. Control: untreated cells. **(E)** Co-staining of Lamin B1 and Emerin in RTN4 KO (left) or CLIMP63 OE (right) cells, as quantified in (D). Arrowheads as in (B). Images: single focal plane confocal images; scale bars: 5 μm. Sample size: N = 226, 278 or N = 365, 328, 82 from left to right in (C); N = 90, 86, 74 or N = 90, 120 from left to right in (D). Quantification: ^∗∗∗^p = 0.0003 (C); ^∗∗∗^p = 0.0002 (control vs RTN4 KO1 in (D)); ^∗∗∗^p = 0.0006 (control vs RTN4 KO2 in (D)); ^∗∗∗∗^p < 0.0001 (control vs CLIMP63 OE in (D)), or not significant (not labelled); two-tailed Fisher’s exact tests.

Strikingly, all three orthogonal methods resulted in NE regions where proteins were mis-localised (**Fig. 3 B**). There were two main types of NE protein mis-localisation events: 1) depletion of non-core NE proteins alongside enrichment of core NE proteins and 2) co-depletion of both core and non-core NE proteins. The core NE protein LAP2α behaved uniquely and only showed enrichment at NE regions with non-core NE protein depletion. 12-18% of nuclei in the converted condition versus 1% of control nuclei had NE regions with protein mis-localisation, as determined using different pairs of core and non-core NE markers (**Fig. 3 C**). Similarly, 16% or 14% of nuclei in two independent RTN4 knockout clones, or 43% of nuclei after CLIMP63 overexpression, had NE regions with protein mis-localisation, as compared to 1% of control nuclei (**Fig. 3, D and E**). The more dramatic effect seen in CLIMP63 overexpression likely points to effects on NE protein localisation beyond converting the NE assembly pattern.

When mechanical stress was applied through collecting the mitotic cells by a shake-off procedure and then re-plating the cells prior to staining (**Fig. 4 A**), we observed a larger fraction of nuclei that had NE regions with protein mis-localisation in all three conditions. Upon mechanical stress, 57% of converted^M^ nuclei (M denotes mechanical stress) versus 8% of control nuclei had NE regions with protein mis-localisation (**Fig. 4 B**). Similarly, 32% or 42% of nuclei in two independent RTN4 knockout clones (**Fig. 4 C**), and 48% of nuclei after CLIMP63 overexpression (**Fig. 4 D**), had NE regions with protein mis-localisation, as compared to 10% of control nuclei. These results suggest that the pre-existing fragility in the NE, which result from promoting NE assembly by lateral sheet expansion in RPE-1 cells, are exacerbated by mechanical stress.

**Figure 4.**
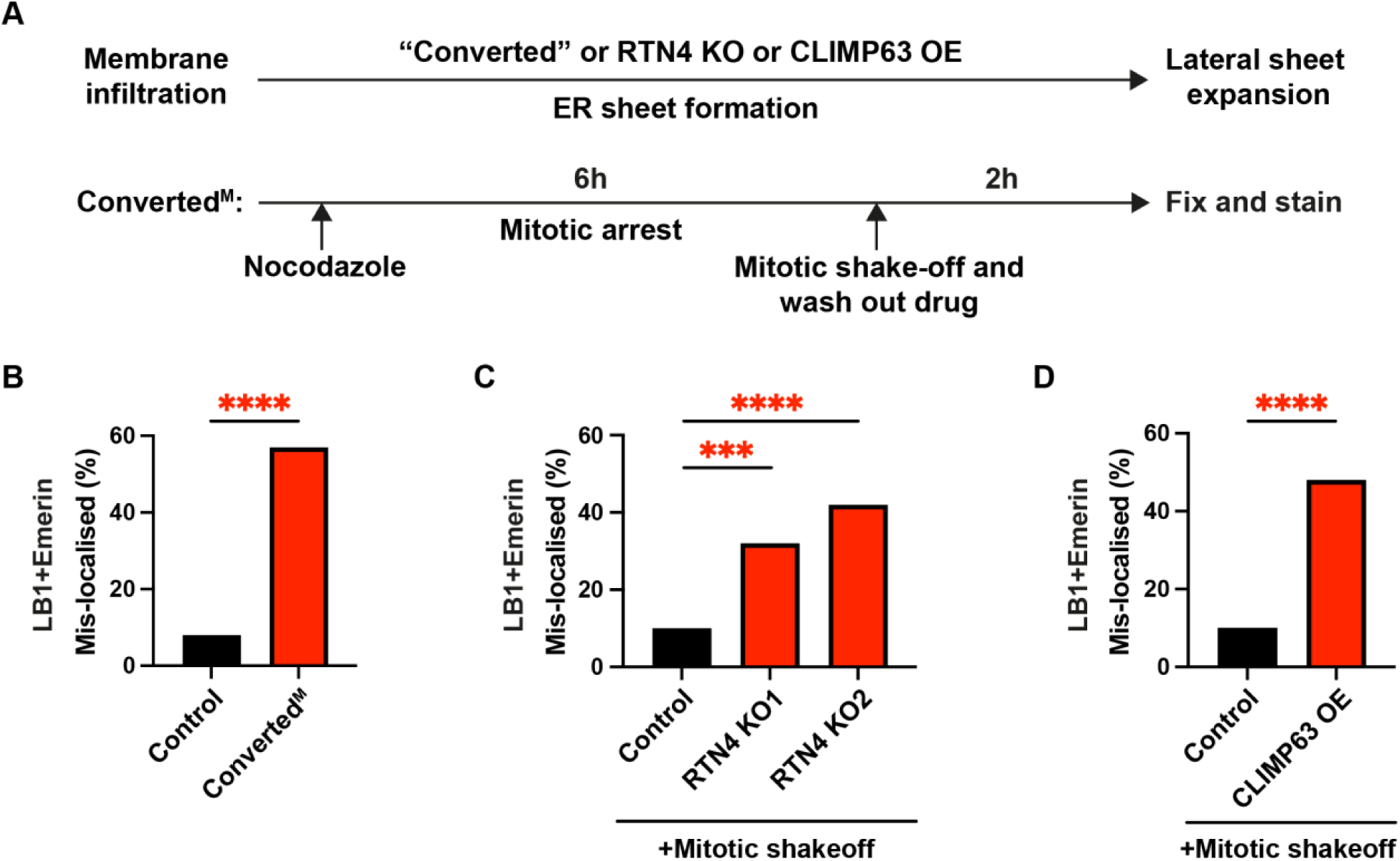
Mechanical stress exacerbates pre-existing NE fragility in RPE-1 cells. **(A)** Experimental scheme as in Fig. 3A except for Converted^M^: depolymerisation of microtubules in interphase and mitosis by Nocodazole-treatment, followed by mitotic shake-off and washing out the drug ((Zhao et al., 2023); Materials and methods). **(B)** Percentage of control or converted^M^ nuclei with NE regions where both Lamin B1 and Emerin (LB1+Emerin) are mis-localised. Control: depolymerisation of microtubules only in mitosis by Nocodazole-treatment, followed by mitotic shake-off and washing out the drug, which does not convert the membrane infiltration NE assembly pattern into lateral sheet expansion ((Zhao et al., 2023); Materials and methods). **(C)** Percentage of nuclei with NE regions where both Lamin B1 and Emerin (LB1+Emerin) are mis-localised in control or two RTN4 KO clones (KO1 and KO2). Control: untreated cells. Note: mitotic shake-off was performed on asynchronous control or RTN4 KO cells. **(D)** Same as in (C) except for CLIMP63 overexpressing (OE) cells. Sample size: N = 428, 556 for control, converted^M^ in (B); N = 77, 223, 88 for control, KO1, KO2 in (C); N = 77, 149 for control, CLIMP63 OE in (D). Quantification: ^∗∗∗∗^p < 0.0001, ^∗∗∗^p < 0.0002 (C), two-tailed Fisher’s exact tests.

In summary, our data show that promoting NE assembly by lateral sheet expansion in normal cells results in NE regions with protein mis-localisation.

### Mis-localised core and non-core proteins mark fragile NEs that are prone to rupture

Core proteins accumulate in the NE during NE rupture repair (Coy et al., 2023; De Vos et al., 2011; Denais et al., 2016; Halfmann et al., 2019; Young et al., 2020). Therefore, we investigated by same-cell correlative live-cell fixed-cell imaging whether the nuclei that have NE regions with protein mis-localisation are prone to rupture. We first followed the formation of nuclei in converted^M^ RPE-1 cells by time-lapse imaging and then stained for various NE markers in the same nuclei to assess their localisation (**Fig. 5 A**). Using GFP-H2B as a live-cell DNA marker, we tracked the formation of individual nuclei for two hours, from prometaphase to early interphase (∼1.5 hours into G1). To exclude any effect of aberrant cell division on nucleus reformation, we excluded multi-polar divisions or chromosome mis-segregation events (that generate lagging chromosomes or chromosomes bridges) from our analysis. In parallel, we used RFP-NLS as a live-cell marker to detect NE ruptures.

**Figure 5.**
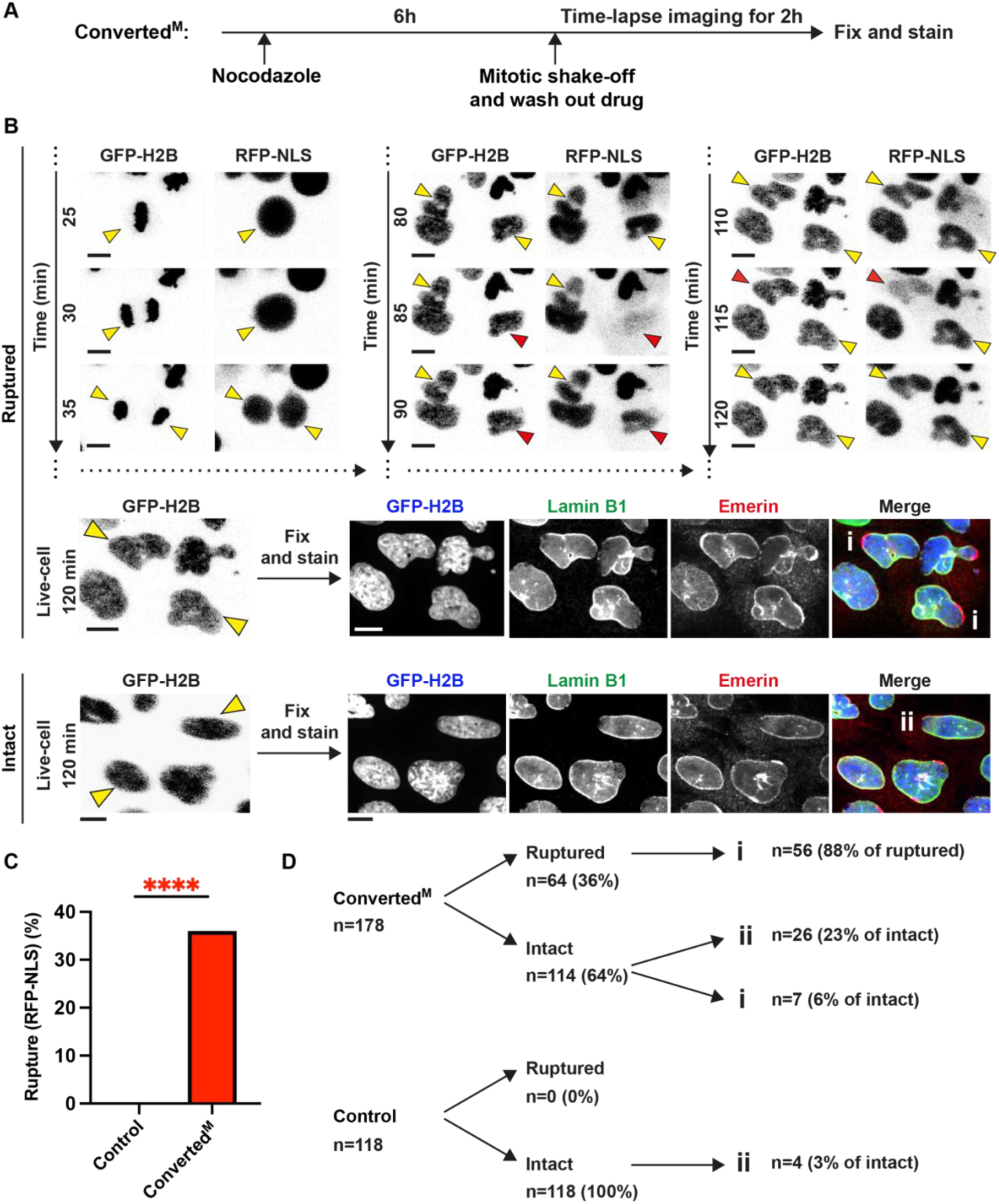
Mis-localised NE proteins mark fragile nuclei that are prone to rupture in converted^M^ RPE-1 cells. **(A)** Experimental scheme of the correlative live-cell fixed cell imaging experiment. Converted^M^ as in Fig. 4A. Nuclei were tracked for 2 hours by time-lapse imaging before the same nuclei were fixed and stained for NE markers. **(B–D)** Correlating NE integrity (RFP-NLS) with NE protein localisation in the same cell. **(B)** Time-lapse images showing converted^M^ nuclei (GFP-H2B) that ruptured transiently in interphase (red arrowheads: nuclear RFP-NLS is lost into the cytoplasm). Yellow arrowheads: the newly assembled nuclei tracked throughout mitosis and early interphase. Note: images from timepoints 0–20 min are not shown; the same time-lapse images of intact nuclei are shown in Fig. S3 B. The following protein mis-localisation events were observed (as marked in the Merge images): (i) ruptured nuclei with NE regions of both Lamin B1 (non-core) depletion and Emerin (core) enrichment; and (ii) intact nuclei with NE regions of Lamin B1 and Emerin co-depletion. **(C)** Percentage of ruptured nuclei (RFP-NLS) in control or converted^M^ cells. Control as in Fig. 4B. **(D)** Percentage of the indicated NE protein mis-localisation events (as marked in the Merge images in (B)) in ruptured or intact nuclei in converted^M^ or control cells. Images: single focal plane widefield (live-cell) or confocal (fixed cell) images; scale bars: 10 μm. Sample size: N = 118, 178 for control, converted^M^ in (C). Quantification: ^∗∗∗∗^p < 0.0001, two-tailed Fisher’s exact test.

Strikingly, the newly formed converted^M^ nuclei underwent extensive NE ruptures, where RFP-NLS was transiently lost from 36% of converted nuclei; by contrast, none of the control nuclei ruptured (**Fig. 5, B and C**). Consistent with prior work, these transient NE ruptures were repaired (Halfmann et al., 2019; Young et al., 2020). 88% of the ruptured nuclei had NE regions both depleted for Lamin B1 (non-core) and enriched for Emerin (core) proteins (**Fig. 5, B and D**), suggesting that non-core protein depletion and core protein enrichment mark ruptured NEs, consistent with prior work (Coy et al., 2023; De Vos et al., 2011; Halfmann et al., 2019; Young et al., 2020). We did not detect Emerin enrichment at the Lamin B1-depleted NE regions in the remaining ruptured nuclei (12%), either because its accumulation was weak or because its accumulation had been resolved (Coy et al., 2023; Halfmann et al., 2019). As expected, the fraction of ruptured nuclei with NE regions of both Lamin B1 depletion and Emerin enrichment increased with mechanical stress (**Fig. S3 A**), consistent with prior studies (Hatch and Hetzer, 2016; Takaki et al., 2017).

Surprisingly, the core and non-core NE proteins also mis-localised in 23% of intact nuclei that did not undergo RFP-NLS loss during time-lapse imaging (**Fig. 5, B and D; and Fig. S3 B**). By contrast to the ruptured nuclei, these intact nuclei had NE regions co-depleted for Lamin B1 and Emerin (**Fig. 5 B**). Given that these NE regions were present in only 3% of intact nuclei in control cells (**Fig. 5 D**), our results suggest that the co-depletion of core and non-core proteins mark fragile NEs that are prone to rupture. A small proportion of the intact nuclei had NE regions with both Lamin B1 depletion and Emerin accumulation (6%) (**Fig. 5 D**). This group represents nuclei where either NLS loss was not detected during time-lapse imaging (five-minute temporal resolution), or the NE ruptured between the end of time-lapse imaging and cell fixation. The remaining intact nuclei (71%) showed no NE defects.

Additionally, we tracked the converted^M^ nuclei for one full cell cycle (from mitosis to mitosis) by time-lapse imaging (**Fig. S4 A**). For this analysis, we used RPE-1 p53 knockout cells to overcome the p53-dependent G1 cell cycle arrest induced by Nocodazole-treatment (Crasta et al., 2012). Most NE ruptures in the converted^M^ nuclei occurred within the first six hours after mitotic exit and decreased with cell cycle progression (**Fig. S4 B**), consistent with the NE fragility originating from NE assembly. Additionally, most NE ruptures were repaired within 5-15 minutes, and the majority of nuclei underwent repeated NE ruptures (**Fig. S4, C and D**). In contrast to previous findings, we did not observe any effect of p53 knockout on NE rupture rates (**Fig. 5 C and Fig. S4 B**; (Yang et al., 2017)).

In summary, promoting NE assembly by lateral sheet expansion in normal cells results in the formation of fragile NEs that are prone to rupture.

## Discussion

Here, using a panel of 13 mammalian cell lines, we demonstrate that the distinct mitotic ER morphologies and NE assembly patterns in normal cells and some cancer cells are located on the opposite ends of the “tubule-sheet continuum”. In all 9 normal mammalian cell lines examined, the mitotic ER is dominated by ER tubules or small sheets and the NE assembles by membrane infiltration. By contrast, in 3 out of the 4 cancer cell lines examined, the mitotic ER is dominated by large ER sheets and the NE assembles by lateral sheet expansion. Thus, based on the cell lines examined, we propose that normal cells cluster on the “ER tubules or small sheets/membrane infiltration” end of the tubule-sheet continuum, whereas cancer cells distribute in between the two opposite ends, and are biased towards the “ER sheets/lateral sheet expansion” end of the continuum. Further, we show that the distinct NE assembly patterns control NE integrity, as promoting NE assembly by lateral sheet expansion in normal cells results in NE fragility. Thus, we propose that NE assembly by lateral sheet expansion is a source of NE fragility in cancer cells.

### The distinct mitotic ER morphologies in normal cells and some cancer cells

A prior study reported similar mitotic ER lengths in NRK-52E (normal) and Huh-7 (cancer) cells (from thin section EM images (Puhka et al., 2012)), which contrasts with our findings showing significantly shorter median ER lengths in mitotic NRK-52E as compared to mitotic Huh-7 cells (**Fig. 1, A and B**). Several findings support our conclusions. First, the density of the mitotic actin cytoskeleton in the subcortical cytoplasm, which is required for maintaining mitotic ER tubules or small sheets (Zhao et al., 2023), is significantly higher in NRK-52E as compared to Huh-7 cells (**Fig. S1**). Second, NRK-52E and Huh-7 cells assemble the NE using distinct patterns that are controlled by mitotic ER morphology: membrane infiltration in NRK-52E and lateral sheet expansion in Huh-7 (**Fig. 2 C**). Together, these data suggest that the mitotic ER is dominated by ER tubules or small sheets in NRK-52E (normal) cells and by large ER sheets in Huh-7 (cancer) cells.

Additionally, fenestrated ER sheets were identified in mitotic SVG-A (normal) and mitotic SUM159 (cancer) cells (Chou et al., 2021). However, the presence of fenestrated ER sheets in few regions of the cell may not reflect the overall mitotic ER morphology. In fact, we previously showed that fenestrated ER sheets are also present in cell types whose overall mitotic ER morphology is dominated by ER tubules or small sheets (Zhao et al., 2023).

### NE fragility from NE assembly by lateral sheet expansion

During lateral sheet expansion, NPCs do not assemble on the NE extensions from large ER sheets, as occurs in the inner-core chromatin regions in HeLa cells (Lu et al., 2011; Otsuka et al., 2016; Zhao et al., 2023). Upon mitotic exit, these NE regions that initially lack NPCs, which are also depleted of other non-core proteins such as Lamin B1 and LBR (Dechat et al., 2004; Haraguchi et al., 2008; Haraguchi et al., 2000; Liu et al., 2018), persist into early interphase as “pore-free islands” and gradually assemble NPCs through the interphase NPC assembly pathway (Doucet et al., 2010; Dultz and Ellenberg, 2010; Maeshima et al., 2006; Mimura et al., 2017; Otsuka et al., 2016; Otsuka and Ellenberg, 2018; Otsuka et al., 2023). One possibility is that core NE proteins become depleted at these pore-free islands, hence forming the fragile NE regions in RPE-1 nuclei after promoting NE assembly by lateral sheet expansion (**Fig. 3 B**). Therefore, we suggest that promoting NE assembly by lateral sheet expansion in normal cells increases the formation of pore-free islands, which are a source of fragile NE regions that are prone to rupture. The gradual assembly of NPCs in pore-free islands through interphase NPC assembly is consistent with our observations that most NE ruptures in converted RPE-1 nuclei occurred within the first 2–6 hours after mitotic exit, and the frequency of NE ruptures decreased with interphase progression (**Fig. S4 B**).

After NE rupture, core proteins accumulate in the NE regions where non-core proteins are depleted, consistent with prior studies (Coy et al., 2023; Denais et al., 2016; Halfmann et al., 2019; Young et al., 2020). Using same-cell correlative live-cell fixed cell imaging, we found that NE proteins were also mis-localised in intact nuclei after the membrane infiltration NE assembly pattern was converted into lateral sheet expansion. These intact nuclei have NE regions depleted for both core and non-core proteins. Thus, we find that the co-depletion of core and non-core proteins mark fragile NEs that are prone to rupture.

Our findings are consistent with prior work showing that normal cells exhibit little or no NE rupture (De Vos et al., 2011; Takaki et al., 2017; Vargas et al., 2012; Yang et al., 2017). By contrast, NE ruptures are more common in cancer cells and are further increased by mechanical stress (Takaki et al., 2017; Vargas et al., 2012). Interestingly, enhancing actomyosin contractility increases NE rupture in cancer cells, but has little effect on nuclear envelope integrity in normal cells (Takaki et al., 2017). We propose that these findings can be explained, at least in part, by the presence of pre-existing NE fragility in the cancer cell nuclei originating from NE assembly by lateral sheet expansion that are exacerbated by the mechanical stress from increased actomyosin contractility.

In summary, by systematically analysing the mitotic ER morphology and NE assembly pattern in different mammalian cell lines, we determined that normal cells and some cancer cells preferentially assemble the NE using distinct NE assembly patterns. Additionally, we show that the NE assembly patterns have important consequences for NE integrity: while both patterns assemble functional NEs, NE assembly by lateral sheet expansion results in increased NE fragility.

## Materials and methods

### Cell culture

Cell lines were cultured at 37°C with 5% CO2 atmosphere. All media were supplemented with 100 IU/ml penicillin and 100 μg/ml streptomycin (Gibco 15140-122 or Sigma-Aldrich P4333). **Human cancer cell lines**: U2OS, HeLa Kyoto (HeLa K), Huh-7, and HCT116 derived cell lines were grown in DMEM+GlutaMAX (Gibco 10569-010) supplemented with 10% FBS (GeminiBio, BenchMark 100-106 or Sigma-Aldrich F7524). **Normal mammalian cell lines**: NRK-52E and NIH-3T3 derived cell lines were grown in DMEM+GlutaMAX supplemented with 10% FBS. **hTERT-immortalized human primary cell lines**: RPE-1 derived cell lines were grown in in DMEM: F12 without phenol red (Gibco 21041-025) supplemented with 10% FBS. Dermal Microvascular Endothelial (DMVE) Cell, Neonatal derived cell lines were grown in Vascular Cell Basal medium supplemented with the Microvascular Endothelial Cell Growth Kit-BBE (ATCC PCS-110-040). HPNE cells were grown in a 3:1 mixture of DMEM+GlutaMAX: Medium M3 Base supplemented with 10% FBS and 10 ng/ml human recombinant EGF (StemCell Technologies Inc. 78006). HUVEC and TeloHAEC derived cell lines were grown in Vascular Cell Basal Medium supplemented with the Vascular Endothelial Cell Growth Kit-VEGF (ATCC PCS-100-041). TIME derived cell lines were grown in Vascular Cell Basal Medium (ATCC PCS-100-030) supplemented with the Microvascular Endothelial Cell Growth Kit-VEGF (ATCC PCS-110-041). ipn02.3 2lambda derived cell lines were grown in DMEM+GlutaMAX supplemented with 10% FBS. All cell lines for this study were monitored for mycoplasma by DNA staining when relevant experiments were performed (using a 60x or 100x objective, see Live-cell microscopy and Immunofluorescence (IF) microscopy below).

### Stable cell generation

Stable cell lines (Sec61β U2OS, Sec61β HeLa, YFP–STIM1 Huh-7, YFP–STIM1 HCT116, YFP–STIM1 NRK-52E, YFP–STIM1 NIH-3T3, YFP–STIM1 DMVE, YFP–STIM1 HUVEC, YFP–STIM1 TeloHAEC, YFP–STIM1 TIME, and YFP–STIM1 ipn02.3 2lambda) were generated by transduction using the lentiviral AcGFP–Sec61β vector or the YFP–STIM1 retrovirus vector (see DNA constructs). YFP–STIM1 F-tractin–mCherry Huh-7 or YFP-STIM1 F-tractin–mCherry NRK-52E was generated by transducing YFP–STIM1 Huh-7 or YFP– STIM1 NRK-52E stable cells using the F-tractin-mCherry lentivirus vector (see DNA constructs). Cells were infected for 16-24 h with virus in the presence of 2 μL polybrene (TR-1003G, Sigma-Aldrich), washed, and allowed to recover before fluorescence-activated cell sorting (FACS). Virus was generated by transfecting HEK293FT cells with the appropriate packaging plasmids (lentivirus: pMD2.G and psPAX2; retrovirus: pGAG/POL and pVSV-G) using Lipofectamine 3000 (Life Technologies) according to the manufacturer’s instructions. The stable cell lines YFP–STIM1 RPE-1, GFP–CLIMP63 RPE-1, and the RTN4 knockout RPE-1 clones (KO1 and KO2) were from (Zhao et al., 2023). The stable cell line H2B–eGFP and TDRFP–NLS RPE-1 was from (Liu et al., 2018). The stable cell line H2B–eGFP and TDRFP–NLS RPE-1 p53 KO was similarly generated from stable RPE-1 p53 KO cells from (Tang et al., 2022).

### DNA constructs

The AcGFP–Sec61β (lentiviral plasmid 154009, from Daniel Gerlich, deposited by Josef Penninger), YFP–STIM1 (retroviral plasmid 19754, from Anjana Rao), and F-tractin–mCherry (lentiviral plasmid 85131, from Tobias Meyer) constructs were obtained from Addgene.

### Antibodies for immunofluorescence

For immunofluorescence, the following primary antibodies were used: mouse Emerin (1:300, Abcam, ab204987), rabbit Emerin (1:150, VWR, 10086-500), mouse Lamin B Receptor (1:300, Sigma Aldrich, SAB1400151), rabbit Lamin B1 (1:1000, Abcam, ab16048), mouse Lamin A (1:1000, Abcam, ab8980), mouse Lap2α (1:300, Abcam, ab66588), rabbit Lap2β (1:200, Bethyl Laboratories, A304-840A), rabbit NUP133 (1:300, Abcam, ab155990), and mouse mAb414 (1:1000, Abcam, ab24609). Secondary antibodies used were goat anti-rabbit Alexa Fluor 568 (1:1000, Thermo Fischer Scientific, A-11036), goat anti-mouse Alexa Fluor 647 (1:1000, Thermo Fischer Scientific, A-21236), and goat anti-rabbit Alexa Fluor 647 (1:1000, Thermo Fischer Scientific, A-21245).

### Fluorescence labelling for live-cell imaging

Cells were plated onto polymer-treated 35-mm imaging μ-dishes (Ibidi 81156) and treated with the following labelling reagents. For labelling DNA, cells were treated with 1 μM of the cell-permeable DNA dye SiR-DNA and 1 μM of the broad-spectrum efflux pump inhibitor verapamil (both from Cytoskeleton Inc. CY-SC007) in cell culture media for approximately 30 min–1 h before imaging. Cells were washed once with PBS, which was then replaced with Imaging solution for live-cell imaging (described under Live-cell microscopy). To label ER membranes, cells were treated with 1 μM of ER-Tracker Red (Life Technologies E34250) for approximately 30 min–1 h in cell culture media before imaging. Cells were washed once with PBS, which was replaced with Imaging solution for live-cell imaging (described under Live-cell microscopy).

### Drug treatments for live-cell imaging

To eliminate cell type differences in the timing of cleavage furrow ingression relative to NE assembly, which can obscure NE recruitment into the inner-core chromatin region, we inhibited cleavage furrow ingression during time-lapse imaging of NE assembly. For inhibition of cleavage furrow ingression, cells were treated with 150 μM of the ROCK inhibitor Y27632 (ATCC ACS3030) in Imaging solution for 10–25 min before cells were mounted onto the microscope for live-cell imaging. Cells were imaged in the presence of the drug (lasting up to 1 h). Inhibiting cleavage furrow ingression does not affect the timing or pattern of NE assembly (Zhao et al., 2023).

### Experiments converting mitotic ER tubules into sheets

For converting interphase and mitotic ER tubules into sheets (the “converted” condition, Fig. 3 and Fig. S3 A), cells were treated with 100 ng/ml of Nocodazole (Millipore Sigma M1404) for 6 h (Zhao et al., 2023). The cells were washed three times with PBS, which was then replaced with fresh media for 2 h before fixation and immunofluorescence staining.

The protocol for the converted^M^ condition is the same as for the converted condition except with additional mechanical stress. Cells were treated with 100 ng/ml of Nocodazole for 6 h, and the arrested cells were collected by a shake-off procedure and washed three times with PBS before replating (Fig. 4, Fig. 5, Fig. S3, and Fig. S4). Mitotic cells were plated onto imaging dishes that were pre-coated with 0.1 mg/ml poly-d-lysine hydrobromide (Sigma Aldrich P6407) to facilitate attachment of mitotic cells. Cells were fixed and stained after 2 h with (Fig. 5) or without (Fig. 4) time-lapse imaging during this time. In the long-term imaging experiments (Fig. S4), cells were followed by time-lapse imaging for one full cell cycle (mitosis to mitosis) after replating.

As controls, cells were treated with Nocodazole in mitosis only, which does not convert mitotic ER tubules into sheets (Zhao et al., 2023). Cells were synchronized in G2/M by treating with 9 μM of the CDK1 inhibitor RO-3306 (Millipore Sigma 217599) for 19 h. Arrested cells were washed five times with PBS and then released into media containing 100 ng/ml of Nocodazole for 6 h. The subsequent steps in the controls were the same as for the converted (washing out the drug) or converted^M^ (mitotic shake-off and washing out the drug) conditions.

For inducing mechanical stress in the RTN4 knockout or CLIMP63 overexpressing RPE-1 cells (Fig. 4), mitotic cells from asynchronous populations were released by mitotic shake-off.

### Live-cell microscopy

The confocal and Airyscan live-cell imaging experiments (Fig. 2 and Fig. S2) were performed on asynchronous cells. Cells were plated onto polymer-treated 35-mm imaging μ-dishes 16-24 h before imaging. Cells were imaged in Imaging solution: HBSS (Gibco) containing 10 mM HEPES (Gibco) and Prolong Live Antifade Reagent (Life Technologies P36974) in the presence of the ROCK inhibitor Y27632 as described above. Each imaging dish was imaged for up to 1 h.

The confocal images were collected on a Nikon Ti-E inverted microscope (Nikon Instruments, Melville, NY) with a Yokogawa CSU-22 spinning disk confocal head with Borealis modification. An In Vivo Scientific stage-top CO2 incubator was used to maintain samples at 37°C. Acquisition parameters, shutters, filter positions and focus were controlled using Metamorph software (Molecular Devices). Laser excitation used: 488-nm, 561-nm, and 642-nm lasers. The images were acquired using a Plan Apo 60x/1.4 NA oil or a Plan Apo 100x/1.45 NA oil objective with a BSI Prime sCMOS camera (Photometrics).

The Airyscan images were collected on an inverted Zeiss LSM 880 microscope equipped with an Airyscan module. Samples were excited using 488-nm, 561-nm, and 633-nm lasers, and imaged with a Plan-Apochromat 63x/1.4NA oil objective. Images were acquired and processed using automatic 2D Airyscan processing in ZEN software (Zeiss).

All images were acquired as z-stacks consisting of 4-6 z-sections with a 1 μm step size. Time-lapse images were acquired at 1-min intervals to capture NE assembly in telophase cells, and image acquisition was stopped when a near-continuous NE was formed, as assessed by eye from the acquired images.

### Correlative live-cell fixed cell imaging experiments

For the live-cell imaging part of the correlative live-cell fixed cell imaging experiments (Fig. 5 and Fig. S3 B), imaging dishes were mounted on a Nikon Ti-E inverted microscope equipped with a Nikon Perfect Focus and a Lumencor LED illumination system. An Okolab cage incubator was used to maintain samples at 37°C and 5% humidified CO_2_. Fluorescence and DIC images of cells expressing H2B–eGFP and RFP–NLS were acquired with five-minute interval using a Plan Apo 20x 0.75 NA air objective with an Andor Zyla 4.2 sCMOS camera. Acquisition parameters, shutters, filter positions, and focus were controlled using Metamorph software (Molecular Devices).

After live-cell imaging, cells were washed with PBS, fixed and processed for immunofluorescence on the same imaging dishes, as described under Immunofluorescence (IF) microscopy. Cells of interest were identified from the recorded movies using the H2B– eGFP channel, and their positions were determined based on the grid pattern on the dishes (Ibidi 81166) using the DIC images. Immunostained samples were then imaged by confocal or Airyscan microscopy (microscope information is described under Live-cell microscopy). 405-nm lasers were used to detect the Hoechst DNA stain, and images were acquired as z-stacks consisting of 8-12 z-sections with a 0.4 μm step size.

### Immunofluorescence (IF) microscopy

Cells were fixed with 4% paraformaldehyde for 15 min, after which cells were washed three times with PBS; cells were then extracted in PBS-0.5% Triton X-100 for 5 min, washed 3 times with PBS, blocked for 30 min in PBS containing 3% BSA (Gemini Bio-Products 700-101P) and incubated with primary antibodies (listed under Antibodies for immunofluorescence) diluted in PBS-3% BSA for 60 min. These samples were then washed 3 times for 5 min with PBS-0.05% Triton X-100. Primary antibodies were detected using species-specific fluorescent secondary antibodies. Cells were incubated with secondary antibodies (listed under Antibodies for immunofluorescence) diluted in PBS-3% BSA for 45-60 min. Finally, the samples were washed 3 times for 5 min with PBS-0.05% Triton X-100 before addition of 2.5 μg/ml Hoechst 33342 to detect DNA. Prolong Diamond Antifade (Life Technologies) was used for mounting of all IF samples.

### Quantification and statistical analysis

#### NE assembly

The ratio of NE recruitment in the inner-core relative to the non-core chromatin region (Fig. 2 C) was calculated from confocal or Airyscan time-lapse images using a custom image analysis pipeline from (Zhao et al., 2023). Briefly, the DNA channel was used to segment the chromosome mass by Li or Otsu fluorescence intensity (FI) thresholding for each timepoint. A 10-pixel wide rim outlining each chromosome mass segmentation was created and mathematically divided into the core (inner-core (IC) and outer-core (OC)) and non-core (NC) chromatin regions. The ER/NE membrane signal (e.g., YFP–STIM1) was extracted using ridge detection (with a sigma of 2) and segmented by FI thresholding. The recruitment of ER membranes to the IC or NC chromatin regions was expressed as the fraction of the segmented region covered by the segmented ER signal. For example: ER recruitment to the IC region = area of the segmented ER signal in the IC region / area of the IC region. The ratio of ER membrane recruitment in the IC relative to the NC region was used to distinguish the two NE assembly patterns. This ratio was calculated at the timepoint of the acquired time-lapse series corresponding to half-maximal ER recruitment for the entire chromosome mass.

#### Mitotic F-actin density in different regions of the cell

The custom image analysis pipeline from (Zhao et al., 2023) was used to measure the density of F-actin in different regions of the cell. The F-tractin–mCherry signal was segmented using ridge detection (with a sigma of 1). The subcortical cytoplasmic region (Fig. S1 B) was defined as the region covered by total ER segmentation minus the region covered by the chromosome rim segmentation. The cell cortex (Fig. S1 B) was defined as a 10-pixel wide rim outlining the region covered by cell segmentation, which was generated using FI thresholding of a cytoplasmic signal (e.g., YFP– STIM1). The density of F-actin in each region was expressed as the fraction of the segmented region covered by the segmented F-tractin–mCherry signal. For example: F-actin density in the subcortical cytoplasm = area of the segmented F-actin signal in the subcortical cytoplasmic region / area of the subcortical cytoplasm.

#### Mitotic ER morphology

Images from the first timepoint of the time-lapse series were used to measure the lengths of anaphase/telophase ER membranes. The ER membrane signal was extracted and segmented using the ImageJ/FIJI plugins “Tubeness” and “Ridge Detection” (Glaser, 2016), as described in (Zhao et al., 2023). For confocal images, membranes shorter than the theoretical resolution limit of diffraction-limited light microscopy (200 nm) were excluded from the analysis. For Airyscan images, membranes shorter than 120 nm were excluded from the analysis. The median ER membrane length was reported for each cell.

#### Scoring NE protein mis-localisation

NE protein mis-localisation was visually determined. A NE was scored as positive for NE protein mis-localisation if the NE contained regions where 1) the non-core NE marker was depleted and the core NE marker was enriched; or 2) the non-core and core NE markers were co-depleted (Fig. 3).

#### Scoring NE ruptures

NE rupture was visually determined. A NE was scored as ruptured if the RFP–NLS signal was lost from the nucleus into the cytoplasm (Fig. 5 and Fig. S4). The onset of rupture was defined as the first timepoint in which the RFP–NLS signal was lost from the nucleus into the cytoplasm. The end of the rupture was defined as the first timepoint where the RFP–NLS signal was fully recovered in the nucleus to pre-rupture levels and there was no detectable RFP–NLS signal in the cytoplasm. The percentage of nuclei that ruptured (Rupture (%) in Fig. 5 C and Fig. S4 B) was measured by counting the number of nuclei that ruptured in the indicated time window, divided by the total number of nuclei examined. A nucleus that underwent multiple ruptures during the time window was counted as a single nucleus. Rupture duration (Fig. S4 C) was defined as the time between the onset and the end of the rupture. The number of ruptures per nucleus (Fig. S4 D) was determined by counting the number of times RFP–NLS was lost after it recovered in the nucleus from a previous rupture.

#### Statistical analysis

All statistical analyses were performed using GraphPad Prism. For the mitotic ER morphology and NE assembly experiments (Fig. 1, Fig. 2, and Fig. S1), non-parametric tests were performed between two (Mann-Whitney) or three or more samples (Kruskal-Wallis test). In the scatter dot plots, bars indicate the mean value of population with 95 % CI. For the NE marker mis-localisation and NE rupture experiments, two-tailed Fisher’s exact tests were performed between two samples (Fig. 3, Fig. 4, Fig. 5, and Fig. S3). No statistical methods were used to predetermine sample size. The experiments were not randomized, and the investigators were not blinded to allocation during experiments and outcome assessment.

## Supplemental materials

Supplemental Table S1 and Supplemental Figures S1-S4, along with the associated table and figure legends, are included at the end of this manuscript.

## Acknowledgments

G. Zhao is supported by the Cambridge Institute for Medical Research (CIMR), and this work was supported by the CIMR Microscopy and Flow Cytometry Core Facilities. This work started in David Pellman’s laboratory at Dana-Farber Cancer Institute and was supported by NIH R37 GM61345-20. The authors would like to thank D. Pellman for the support to independently pursue this work; S. Liu for preliminary experiments; and S. Liu, E. Sydir, and D. Pellman for critical reading of the manuscript.

## Author contributions

G. Zhao conceived the project, designed the experiments, and supervised the study. S. Arun and G. Zhao performed the experiments. G. Zhao analysed the data and wrote the manuscript.

---

**Table S1.** Summary of the 13 mammalian cell lines in this study. The cell type, tissue, disease, or species information of the cell lines. The hTERT-immortalized human primary, normal mammalian, and human cancer cell lines are highlighted in green, blue, and orange respectively. This colour scheme is used throughout the figures.

| Cell line | Cell type | Tissue | Disease | Species |
| --- | --- | --- | --- | --- |
| **hTERT-immortalised human primary** |  |  |  |  |
| RPE-1 | Epithelial cell | Eye; pigmented epithelium; retina | Normal | Human |
| Dermal microvascular endothelial cell (DMVE) | Endothelial cell | Skin; dermis | Normal | Human |
| HPNE | Epithelial cell | Pancreas; duct | Normal | Human |
| HUVEC | Endothelial cell | Umbilical cord; vascular endothelium | Normal | Human |
| TeloHAEC | Endothelial cell | Heart; aorta | Normal | Human |
| TIME | Endothelial cell | Skin; dermal vascular endothelium; foreskin | Normal | Human |
| ipn02.3 2lambda | Shwann cell | Sural nerve | Normal | Human |
| **Normal mammalian** |  |  |  |  |
| NRK-52E | Epithelial cell | Kidney | Normal | Rat |
| NIH-3T3 | Fibroblast | Embryo | Normal | Mouse |
| **Human cancer** |  |  |  |  |
| U2OS | Epithelial cell | Bone | Osteosarcoma | Human |
| HeLa | Epithelial cell | Uterus; cervix | Adenocarcinoma | Human |
| Huh-7 | Epithelial cell | Liver | Hepatocellular carcinoma | Human |
| HCT116 | Epithelial cell | Large intestine; colon | Colorectal carcinoma | Human |

**Figure S1.**
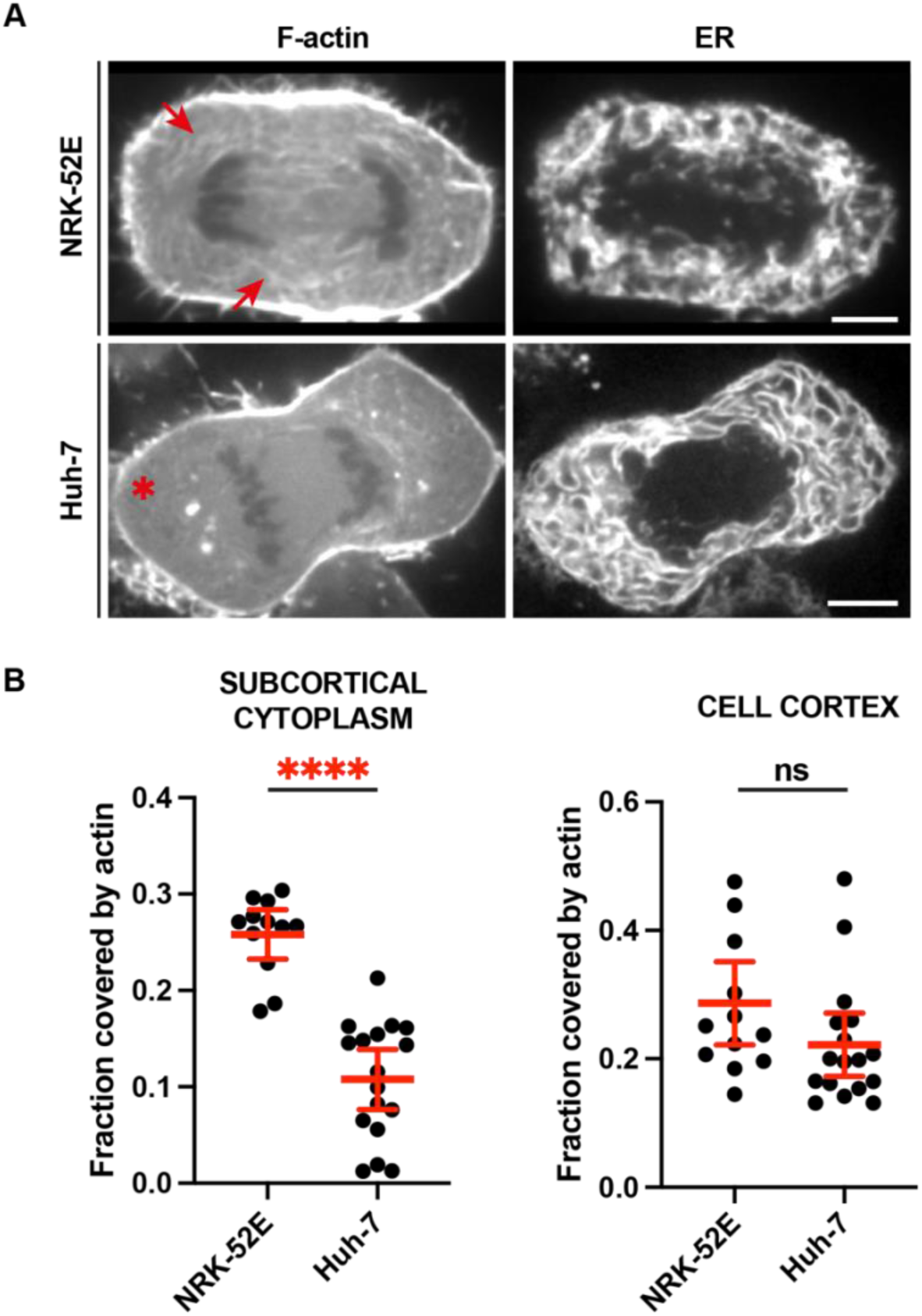
Comparison of F-actin density in anaphase/telophase NRK-52E and Huh-7 cells. **(A)** Confocal live-cell images of the mitotic F-actin (F-tractin-mCherry) or the ER (YFP-STIM1). Note the mitotic F-actin density in the subcortical cytoplasm correlates with mitotic ER morphology: high F-actin density (arrows) and ER tubules/small sheets in NRK-52E cells; low F-actin density (asterisk) and large ER sheets in Huh-7 cells. The median ER lengths in anaphase/telophase NRK-52E or Huh-7 cells are shown in Fig. 1 B. **(B)** The density of mitotic F-actin in the subcortical cytoplasm (left) or cell cortex (right). Note: the higher density of F-actin in the subcortical cytoplasm, but not in the cell cortex, in NRK-52E as compared to Huh-7 cells. Images: single focal plane; scale bars: 5 μm. Sample size: N = 12, 17 for NRK-52E, Huh-7 in (B). Quantification: ^∗∗∗∗^p < 0.0001, or not significant (ns), two-tailed Mann-Whitney tests.

**Figure S2.**
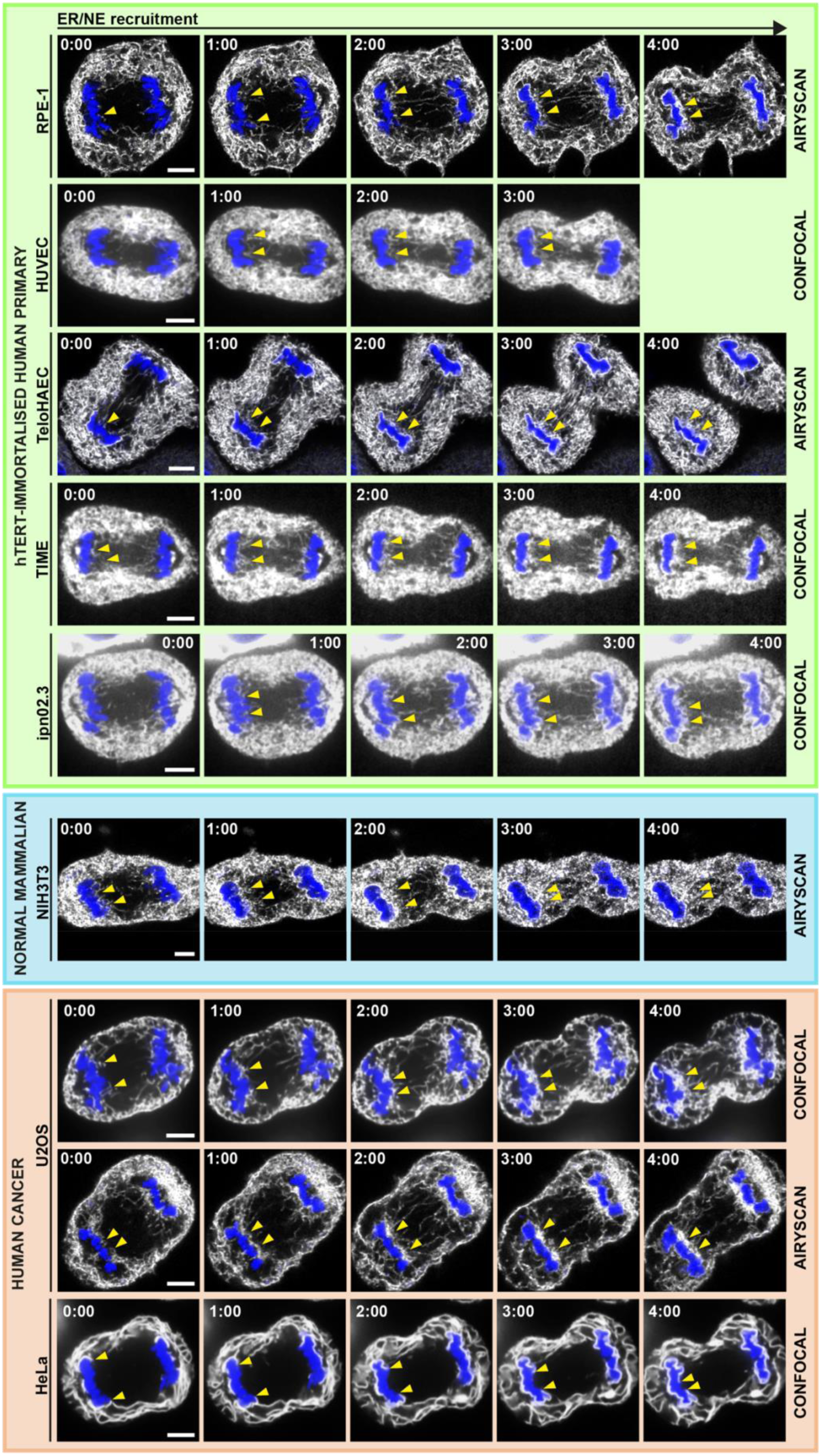
Time-lapse images of NE assembly in additional cell lines (not shown in Fig. 2A) Confocal or Airyscan images (as indicated) from time-lapse series showing NE assembly in additional cell lines quantified in Fig. 2 C. Arrowheads: NE membrane extension into the inner-core (IC) from the non-core (NC) chromatin region by lateral sheet expansion in HeLa cells; or NE recruitment into the IC chromatin region by membrane infiltration in all the other cell lines. Images: single focal plane; scale bars: 5 μm.

**Figure S3.**
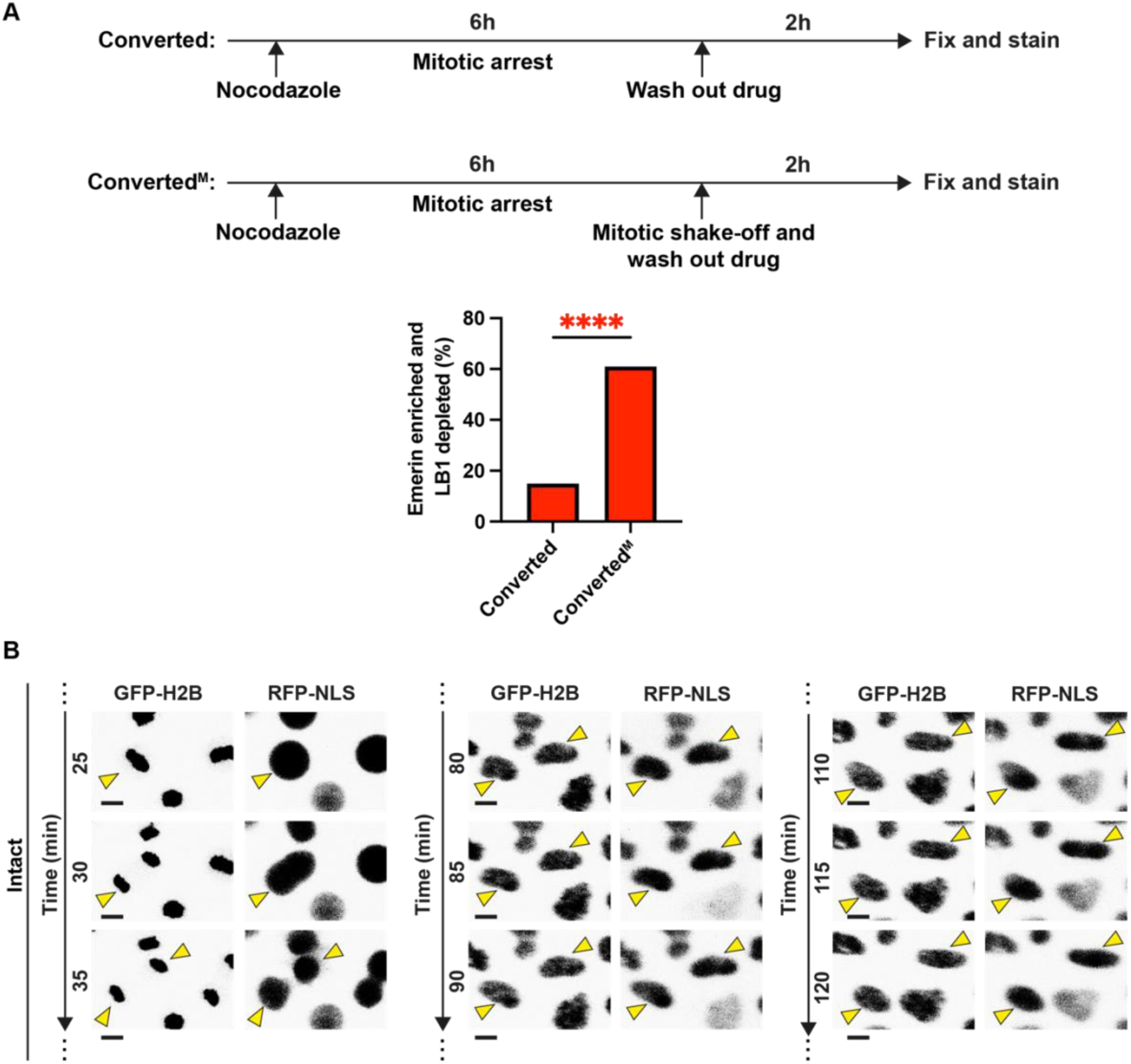
Frequency of ruptured nuclei in converted or converted^M^ RPE-1 cells, and time-lapse images of intact converted^M^ nuclei. **(A)** Top, experimental scheme. Converted (as in Fig. 3A) or converted^M^ (as in Fig. 4A). Bottom, percentage of ruptured nuclei with NE regions of both Lamin B1 depletion and Emerin enrichment. **(B)** Time-lapse images showing converted^M^ nuclei (GFP-H2B) that remained intact throughout the 2 hours of imaging (no RFP-NLS loss from the nucleus into the cytoplasm). The same nuclei were fixed and stained for NE markers as shown in Fig. 5 B. Sample size: N = 278, 556 for converted, converted^M^ in (A). Quantification: ^∗∗∗∗^p < 0.0001, two-tailed Fisher’s exact test.

**Figure S4.**
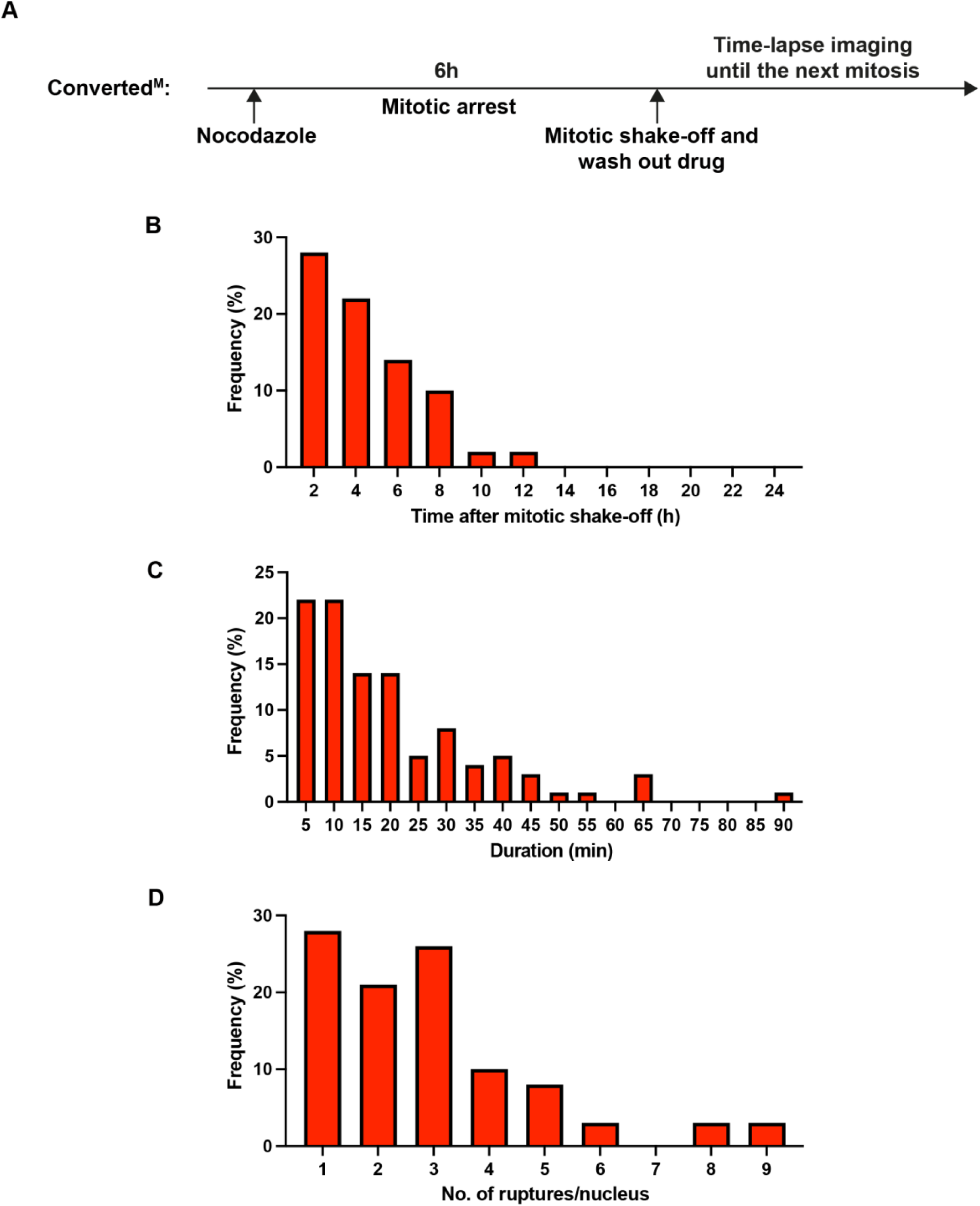
NE rupture timing, duration, and frequency per nucleus in converted^M^ RPE-1 p53 knockout cells in one full cell cycle (mitosis to mitosis) **(A)** Experimental scheme. Converted^M^ as in Fig. 4A. **(B)** Timing (in hours) of NE ruptures (RFP-NLS loss from the nucleus into the cytoplasm) after mitotic shake-off. Shown is the timing of the first NE rupture for each ruptured nucleus. **(C)** Duration of NE ruptures (in minutes) before they were repaired (RFP-NLS recovered into the nucleus). Shown are all NE rupture events: repeat ruptures of the same nucleus are counted as distinct rupture events in this analysis. **(D)** Number of ruptures per nucleus. Sample size: N = 39 nuclei.

## Notes

### Competing Interest Statement

The authors have declared no competing interest.

